# Cyclase-associated protein is a pro-formin anti-capping processive depolymerase of actin barbed and pointed ends

**DOI:** 10.1101/2023.11.30.569482

**Authors:** Ekram M. Towsif, Shashank Shekhar

## Abstract

Cellular actin networks display distinct assembly and disassembly dynamics resulting from multicomponent reactions occurring primarily at the two ends and the sides of actin filaments [1–3]. While barbed ends are considered the hotspot of actin assembly [4], disassembly is thought to primarily occur via reactions on filament sides and pointed ends [3, 5–11]. Cyclase-associated protein (CAP) has emerged as the main protagonist of actin disassembly and remodeling – it collaborates with cofilin to increase pointed-end depolymerization by 300-fold [6, 7], promotes filament “coalescence” in presence of Abp1 [12], and accelerates nucleotide exchange to regenerate monomers for new rounds of assembly [13–15]. CAP has also been reported to enhance cofilin-mediated severing [16, 17], but these claims have since been challenged [7]. Using microfluidics-assisted three-color single-molecule imaging, we now reveal that CAP also has important functions at filament barbed ends. We reveal that CAP is a processive barbed-end depolymerase capable of tracking both ends of the filament. Each CAP binding event leads to removal of about 5,175 and 620 subunits from the barbed and pointed ends respectively. We find that the WH2 domain is essential, and the CARP domain is dispensable for barbed-end depolymerization. We show that CAP co-localizes with barbed-end bound formin and capping protein, in the process increasing residence time of formin by 10-fold and promoting dissociation of CP by 4-fold. Our barbed-end observations combined with previously reported activities of CAP at pointed ends and sides, firmly establish CAP as a key player in actin dynamics.

## RESULTS AND DISCUSSION

### CAP is a barbed-end depolymerase

We employed microfluidics-assisted total internal reflection fluorescence microscopy (mf-TIRF) to investigate effects of mouse CAP1 (referred to as CAP henceforth) on barbed-end depolymerization. Actin filaments with free barbed ends were elongated from coverslip-anchored spectrin-actin seeds by flowing in a solution containing Alexa-488 labelled G-actin and unlabeled profilin (profilin-actin) (Fig. 1A). The filaments were aged for 15 minutes to allow P_i_ release and conversion to ADP-F-actin following which they were exposed to TIRF buffer in absence or presence of CAP. While control filaments depolymerized at 9.7±1.3 subunits/s (su/s hereafter) (± sd), addition of 1 µM CAP caused a 4-fold increase in depolymerization rate (38.2 ± 5.2 su/s) (Fig. 1C – E). Acceleration of barbed-end depolymerization by CAP is consistent with results from a recent study from the Goode lab [18]. Our observation of barbed-end depolymerization together with previous reports of 4-fold acceleration of depolymerization at pointed-ends [6, 13] establish CAP as a bona-fide depolymerase of both ends of actin filaments.

**Fig. 1:**
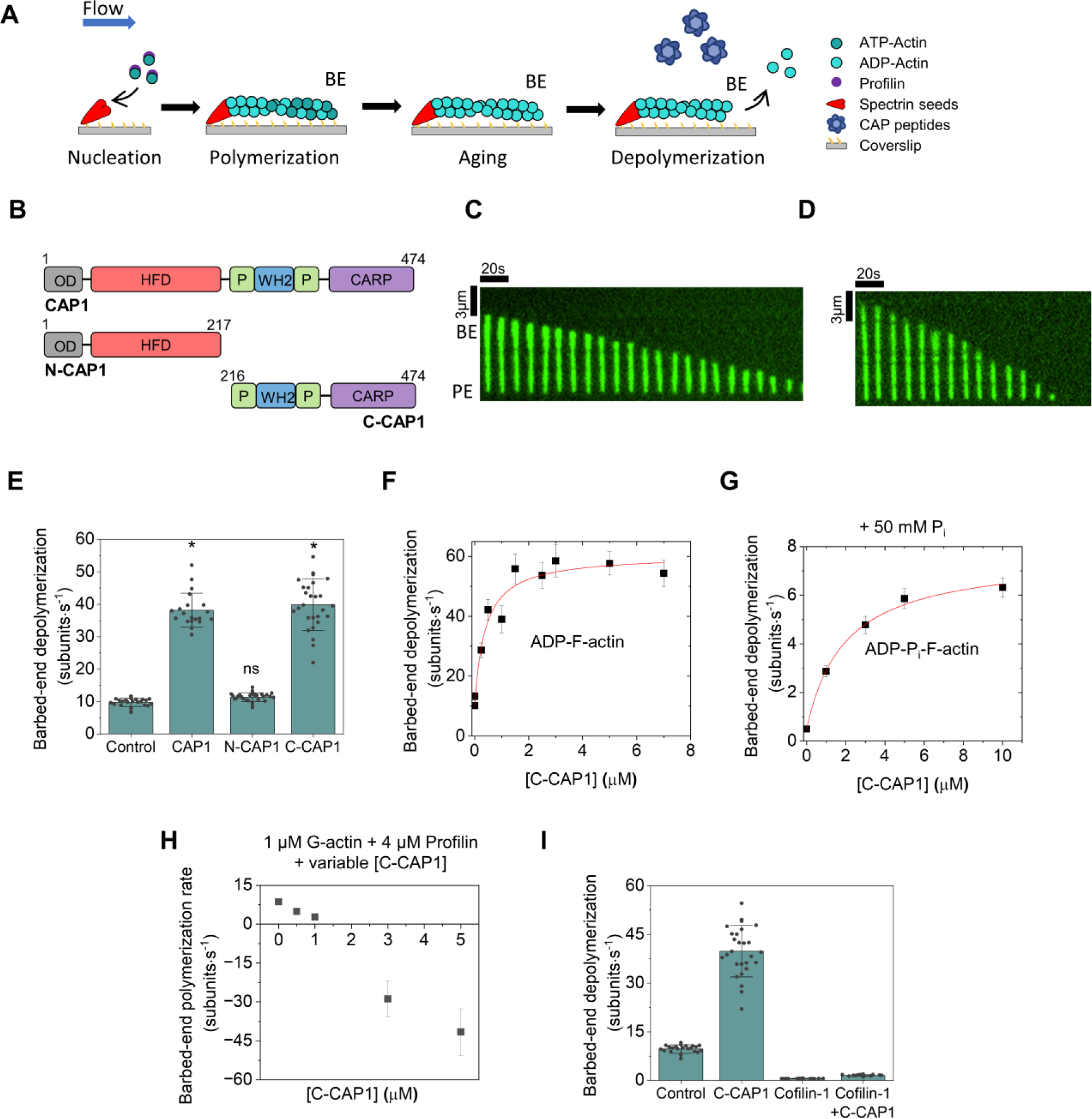
Mouse CAP1 accelerates actin filament barbed-end depolymerization. **(A)** Schematic representation of the experimental strategy. Actin filaments with free barbed ends were polymerized by exposing coverslip-anchored spectrin-actin seeds to 1 µM G-actin (15% Alexa-488 labeled) and 4 µM profilin. The filaments were then aged for 15 minutes and then exposed to 1 µM CAP (full length, N-CAP or C-CAP) or control buffer. Barbed-end depolymerization was monitored over time. BE, barbed end; **(B)** Domain diagram of mouse CAP1 constructs used in this study. **(C)** Representative kymograph of an Alexa-488-labeled actin filament depolymerizing in control buffer. BE, barbed end; PE, pointed end. **(D)** Same as (C) but in the presence of 1 μM CAP1. **(E)** Rates (± sd) of barbed-end depolymerization of ADP filaments in the presence of buffer (control), 1 µM CAP1, 1 µM N-CAP1 or 1 µM C-CAP1. ✲, statistical comparison by two-sample t-test against control (*p* < 0.05). ns, no evidence for significance at *p* = 0.05. Number of filaments analyzed for each condition (left to right): 25, 20, 30 and 29. **(F)** Rates (± sd) of barbed-end depolymerization of ADP filaments as a function of C-CAP1 concentration. *N* = 21 filaments analyzed per concentration. The line is a fit to a hyperbolic binding curve. **(G)** Rates (± sd) of barbed-end depolymerization of ADP-P_i_ filaments as a function of C-CAP1 concentration (same experimental strategy as in panel A was followed with the exception that filaments were maintained in 50 mM P_i_ throughout, see methods). *N* = 20 filaments analyzed per concentration. The line is a fit to a hyperbolic binding curve. **(H)** Rates (± sd) of free barbed-end elongation in the presence of 1 µM G-actin, 4 µM profilin, and different concentrations of C-CAP1. Number of filaments analyzed for each condition (left to right): 20, 20, 20, 17, and 20. **(I)** Rates (± sd) of barbed-end depolymerization of ADP filaments in the presence of 1 µM C-CAP1 and 5 µM cofilin-1 (alone and together). Number of filaments analyzed for each condition (left to right): 25, 29, 20 and 20.

### Barbed-end depolymerization mediated by C-terminal half of CAP

CAP is a modular protein whose N-terminal (N-CAP) and C-terminal (C-CAP) halves function autonomously in actin disassembly and monomer recycling (Fig. 1B) [19–21]. N-CAP consists of an oligomerization domain (OD), a helical-folded domain (HFD), and promotes pointed-end depolymerization [6, 7]. N-CAP has also been reported to bind filament sides and enhance cofilin-mediated severing [16, 17]. However, these conclusions have been challenged by a recent structural study which implied that actin-binding HFD domains of N-CAP could only be docked at the filament pointed end [7]. Consistently, the authors observed no enhancement of cofilin-mediated severing by N-CAP. C-terminal half of CAP or C-CAP consists of WH2 domain [22], CARP domain and polyproline motifs. WH2 and CARP domains bind G-actin, accelerate nucleotide exchange (from ADP to ATP) to regenerate ATP-actin monomers for new rounds of polymerization [13–15]. Lastly, while N-CAP monomers oligomerize into tetramers [23, 24] and/or hexamers [16, 17], C-CAP monomers self-assemble into dimers [15, 25].

In light of previously reported depolymerization function of N-CAP, we wondered if it might also be responsible for barbed-end depolymerization by CAP. Free barbed ends of ADP-actin filaments were exposed to a solution containing either N-CAP or C-CAP. While N-CAP had no effect on barbed-ends, C-CAP accelerated their depolymerization by about 4-fold, similar to full-length CAP (Fig. 1E). Thus, the depolymerization action of CAP at the two ends is facilitated by its distinct halves i.e., N-CAP at the pointed and C-CAP at the barbed end. Additional analysis showed that C-CAP promotes depolymerization in a concentration-dependent fashion (Fig. 1F). At saturation, a maximal depolymerization rate of about 58 su/s, approximately 6-fold higher than control, was observed. These kinetics suggest that at lower C-CAP concentrations, binding of C-CAP to actin filaments might be rate-limiting.

### Effects of filament age, cofilin and polymerizable G-actin

While actin networks *in vivo* can turn over in just a few seconds [26–29], depolymerization *in vitro* is orders of magnitude slower, mainly due to slow P_i_ release [30, 31]. We therefore wondered if C-CAP might also depolymerize unaged filaments. To prevent aging, actin filaments were maintained in TIRF buffer containing 50 mM P_i_, as done previously [32]. We found that C-CAP also accelerated barbed-end depolymerization of ADP-P_i_ filaments (Fig. 1G). Notably, although average depolymerization rate of ADP-P_i_ filaments by C-CAP was 10-fold lower than that of ADP filaments (∼6 vs ∼60 su/s), the fold-increase over control was almost double for ADP-P_i_ than ADP filaments (12-fold vs 6-fold). Thus, C-CAP is capable of accelerating depolymerization of both aged and newly-assembled actin filaments.

While our experiments thus far were conducted in absence of G-actin, cytosol contains high amounts of assembly-competent actin monomers, majority of which are bound to profilin [2, 33, 34]. We therefore investigated how presence of profilin-actin monomers affected depolymerization by C-CAP. In the presence of 1 µM profilin-actin, filament barbed ends elongated at 8.6 ± 0.5 su/s (Fig. 1H). Further addition of C-CAP significantly altered this behavior. While low concentrations of C-CAP slowed net polymerization, higher concentrations induced net depolymerization. Addition of 1 µM profilin-actin to reactions containing 5 µM C-CAP led to a 30% reduction in net depolymerization rate from 57.7 ± 3.9 su/s to 41.8 ± 8.9 su/s. It is unclear whether this reduction results from sequestration of actin monomers by C-CAP or due to C-CAP’s barbed end interaction. Notably, a known barbed-end depolymerase, twinfilin, also shows a similar behavior, both in absence (for ADP-P_i_ filaments) and presence of polymerizable actin monomers [32].

We then asked if C-CAP, like N-CAP, might also synergize with cofilin [6, 7]. Supplementing C-CAP only reactions with cofilin (human Cofilin-1) almost completely extinguished barbed end depolymerization (Fig. 1I), leading to rates only slightly higher than cofilin-only reactions. Indeed, a previous study reported that cofilin’s saturation of filament sides drastically reduces barbed-end depolymerization [35]. Our data suggests that C-CAP is not able to catalyze disassembly of cofilin-coated filaments. Cofilin thus has opposing effects on CAP’s depolymerization abilities at the two ends of actin filaments.

### Barbed-end depolymerization by CAP is conserved and requires WH2 domains

CAP is a highly conserved protein expressed across fungus, plants and vertebrates [3]. We therefore asked if barbed-end depolymerization by CAP was conserved. Similar to the mammalian homolog, the C-terminal half of *S. cerevisiae* Srv2 (C-Srv2) also accelerated depolymerization. Notwithstanding the qualitative similarity, even at high concentration (∼5 µM) C-Srv2 only caused a 1.8-fold increase in depolymerization as compared to the 6-fold increase by C-CAP (Fig. 2A-C). Like N-CAP, N-Srv2 had no impact on barbed end depolymerization. Thus, combined with our previous pointed-end studies [6], our results here show that Srv2/CAP’s promotion of depolymerization at both filament ends is conserved between budding yeast and human homologs.

**Fig. 2:**
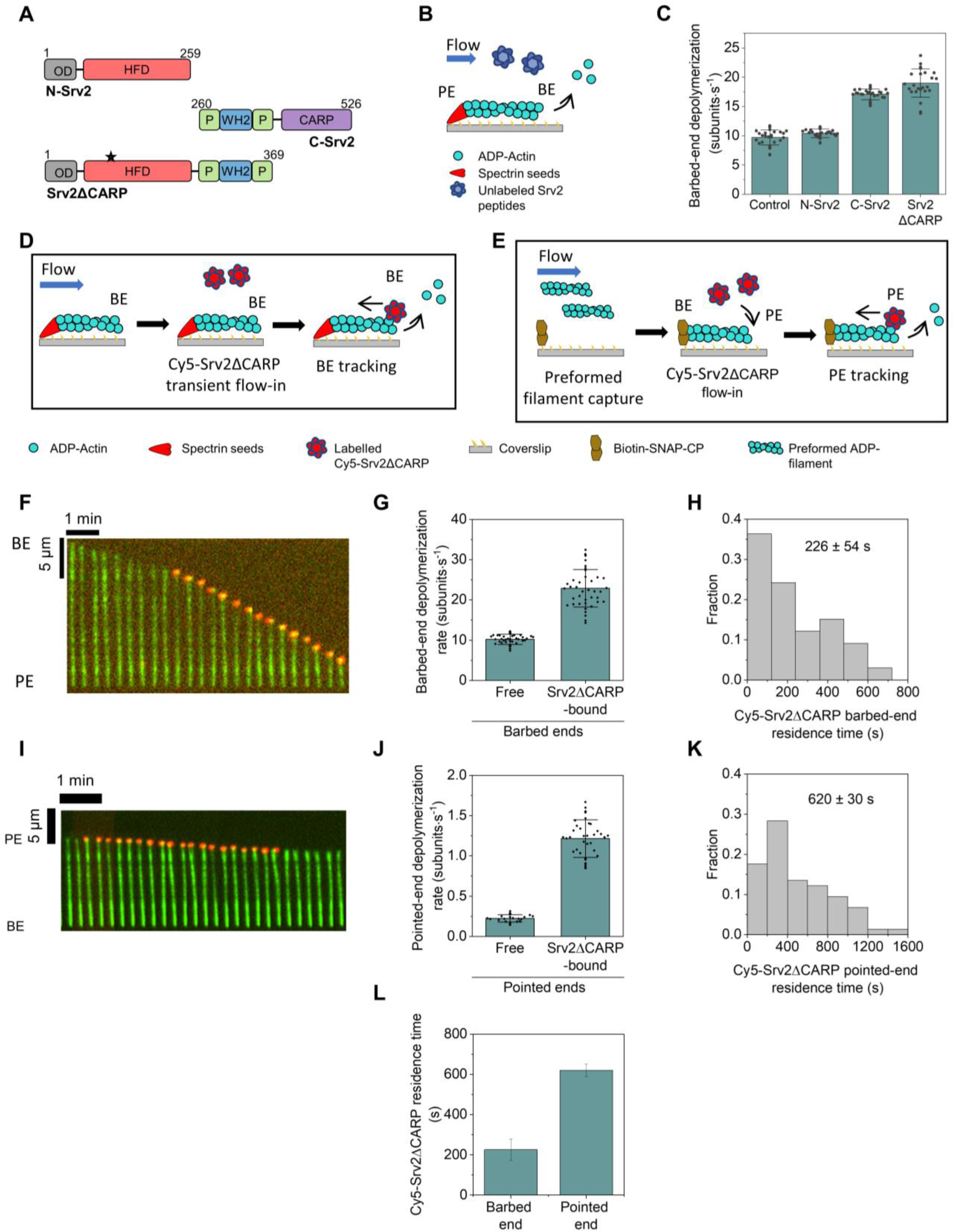
Srv2/CAP is an evolutionary conserved processive depolymerase of both ends of actin filaments. **(A)** Domain diagram of *S. cerevisae* Srv2/CAP constructs used in this study. Black asterisk on Srv2ΔCARP indicates the location of its single cysteine, which was used for dye-labeling. **(B)** Schematic representation of the experimental strategy. ADP-actin filaments with free barbed ends were assembled similarly as in Fig. 1A and exposed to the described Srv2 constructs. Blue colored Srv2 molecules represent unlabeled protein. **(C)** Rates (± sd) of barbed-end depolymerization of ADP filaments in the presence of buffer (control), 5 µM N-Srv2, 5 µM C-Srv2 and 5 µM Srv2ΔCARP. **(D)** Schematic representation of the experimental strategy for single-molecule imaging for visualizing Cy5-Srv2ΔCARP at barbed ends. Red colored Srv2ΔCARP molecules represent labeled protein. **(E)** Schematic representation of the experimental strategy for single-molecule imaging for visualizing Cy5-Srv2ΔCARP at pointed ends. **(F)** Representative kymograph of an Alexa-488-labeled actin filament (green) with Cy5-Srv2ΔCARP (red) processively tracking its barbed end. **(G)** Rates (± sd) of barbed-end depolymerization of free and Cy5-Srv2ΔCARP-bound barbed ends in buffer (no free Cy5-Srv2ΔCARP in solution). Number of filaments analyzed for each condition (left to right): 44 and 42. **(H)** Distribution of lifetimes of Cy5-Srv2ΔCARP at barbed ends (n= 33 Cy5-Srv2ΔCARP-bound ends). **(I)** Representative kymograph of an Alexa-488-labeled actin filament (green) with Cy5-Srv2ΔCARP (red) processively tracking filament pointed end. **(J)** Rates (± sd) of barbed-end depolymerization of free and Cy5-Srv2ΔCARP-bound pointed ends in buffer (no free Cy5-Srv2ΔCARP in solution). Number of filaments analyzed for each condition (left to right): 25 and 40. **(K)** Distribution of lifetimes of Cy5-Srv2ΔCARP at pointed ends (n= 74 Cy5-Srv2ΔCARP-bound ends) **(L)** Average lifetime of Cy5-Srv2ΔCARP at barbed and pointed ends.

We then sought to determine which of the two actin-binding domains in the C-terminal half i.e., WH2 or CARP, caused barbed-end depolymerization. We expressed and purified Srv2ΔCARP (Fig. 2A), a new construct which contained the WH2 domain but not the CARP domain. In contrast with a recent study [18], we found that absence of CARP domain did not abolish barbed-end depolymerization (Fig. 2C). Instead, we observed that Srv2ΔCARP and C-Srv2 promoted barbed end depolymerization equally well. Since the only actin binding motif common between the two constructs is the WH2 domain, our results imply that the WH2 domain is necessary, and CARP is dispensable for barbed-end depolymerization by Srv2/CAP.

### Srv2/CAP molecules processively track depolymerizing barbed ends

To further reveal the molecular mechanism by which Srv2/CAP induces barbed-end depolymerization, we directly visualized Cy5-labelled Srv2ΔCARP on actin filaments using single-molecule imaging. We decided to use Srv2ΔCARP for single molecule experiments as Srv2ΔCARP (monomers) contains only a single cysteine which can be fluorescently labelled in a residue-specific manner (Fig. 2C). In contrast, while CAP, C-CAP, and CAP1ΔCARP monomers contain six, four and two, cysteines; Srv2 and C-Srv2 monomers contain 4 and 3 cysteines respectively. In addition, we have previously successfully used Cy5-Srv2ΔCARP to visualize Srv2/CAP molecules on pointed-ends of cofilin-decorated filaments [6].

Free barbed ends of ADP-actin filaments were first transiently exposed to a flow containing Cy5-Srv2ΔCARP (100 nM monomers) in the mf-TIRF chamber, followed by continuous exposure to TIRF buffer (Fig. 2D). We observed that Cy5-Srv2ΔCARP molecules associated directly with actin filament barbed ends. All ends depolymerizing with a detectable Cy5 signal underwent rapid depolymerization. Consistently, the absence of Cy5 signal at the barbed end was accompanied by slow depolymerization and arrival of Cy5 signal was accompanied by initiation of rapid depolymerization (Fig. 2F,G). The depolymerization rate of Cy5-Srv2ΔCARP-bound barbed ends was similar to maximal depolymerization seen at saturating concentrations of unlabeled Srv2ΔCARP (Fig. 2C), suggesting that at saturating Srv2ΔCARP, the filament end is almost continuously occupied by a Srv2ΔCARP molecule. Each Cy5-Srv2ΔCARP binding event on average lasted 226 ± 54 seconds (Fig. 2H) and barbed ends with a visible Cy5 signal depolymerized at 22.9 ± 4.6 su/s (Fig. 2G,H). Product of these two values yields the average number of subunits removed per Srv2ΔCARP binding event, 5,175 ± 1,615 subunits.

Our single-molecule experiments imply that Srv2/CAP is a processive depolymerase of filament barbed ends. So far, *S. cerevisiae* twinfilin is the only other protein reported to processively depolymerize barbed ends [36]. Twinfilin’s processive behavior has however been challenged in light of recent structural insights which implied that a twinfilin molecule might instead only cause dissociation of terminal actin subunits at the barbed end [37].

### Srv2/CAP molecules processively track depolymerizing pointed ends

Two previous studies directly visualized fluorescently labelled Srv2/CAP molecules associating with pointed ends of cofilin-coated filaments [6, 7]. However, due to the transient nature of Srv2/CAP associations (∼2 seconds) with pointed ends of cofilin-actin filaments, it was not possible to ascertain whether Srv2/CAP molecules translocated with depolymerizing pointed ends i.e., were processive. To address this question, we performed single-molecule imaging using Cy5-Srv2ΔCARP. Preformed fluorescent ADP-actin filaments were captured by coverslip-anchored capping protein in a mf-TIRF chamber. Upon a transient exposure to Cy5-Srv2ΔCARP (100 nM), filaments were exposed continuously to TIRF buffer (Fig. 2E). We observed Cy5-Srv2ΔCARP associating with filament pointed ends. All ends with a detectable Cy5 signal underwent rapid depolymerization (Fig. 2I,J). Appearance of Cy5-Srv2ΔCARP signal was accompanied by beginning of rapid depolymerization. Consistently, the disappearance of Cy5-Srv2ΔCARP was accompanied by resumption of slow depolymerization characteristic of free pointed ends depolymerizing in buffer (Fig. 2I). Each binding event on average lasted 620 ± 30 seconds (Fig. 2K, L). Pointed-ends with Cy5 signal depolymerized at 1.2 ± 0.2 su/s (Fig. 2J,K), about 6-fold faster than filaments without a visible Cy5 signal. Product of these two values yields the average number of subunits removed per Srv2ΔCARP binding event, 743 ± 130 subunits. Our data thus supports the view that Srv2/CAP is a processive depolymerase of both ends of actin filaments.

### Srv2/CAP associates with formin-bound barbed ends and slows formin’s dissociation without affecting elongation rate

In light of direct binding of Srv2/CAP to barbed ends, we asked if Srv2/CAP might also associate with barbed ends bound to other ligands such as formin and capping protein, which either accelerate or arrest barbed assembly. We first studied formin. SNAP-tagged formin mDia1 (FH1-FH2-C) was expressed and labeled with benzylguanine functionalized fluorescent dye (649-mDia1). SNAP-tagging and labelling did not alter dimerization or actin assembly properties of formin, as seen previously [38–41].

Alexa-488 labeled actin filaments with free barbed ends were transiently exposed to either 649-mDia1 alone or together with Cy5-Srv2ΔCARP, followed by a solution containing profilin-actin (Fig. 3A). In absence of Cy5-Srv2ΔCARP, the majority of filaments only displayed 649-mDia1 at their barbed end (Fig. 3B). When 649-mDia1 and Cy5-Srv2ΔCARP were simultaneously present, two distinct barbed-end populations were seen : 1) bound only to 649-mDia1 2) bound to both 649-mDia1 and Cy5-Srv2ΔCARP with the two molecules together tracking the elongating end (Fig. 3C). Importantly, the barbed-end elongation rate remained unchanged independent of whether formin was bound alone to barbed ends or together with Cy5-Srv2ΔCARP (Fig. 3D).

**Fig. 3:**
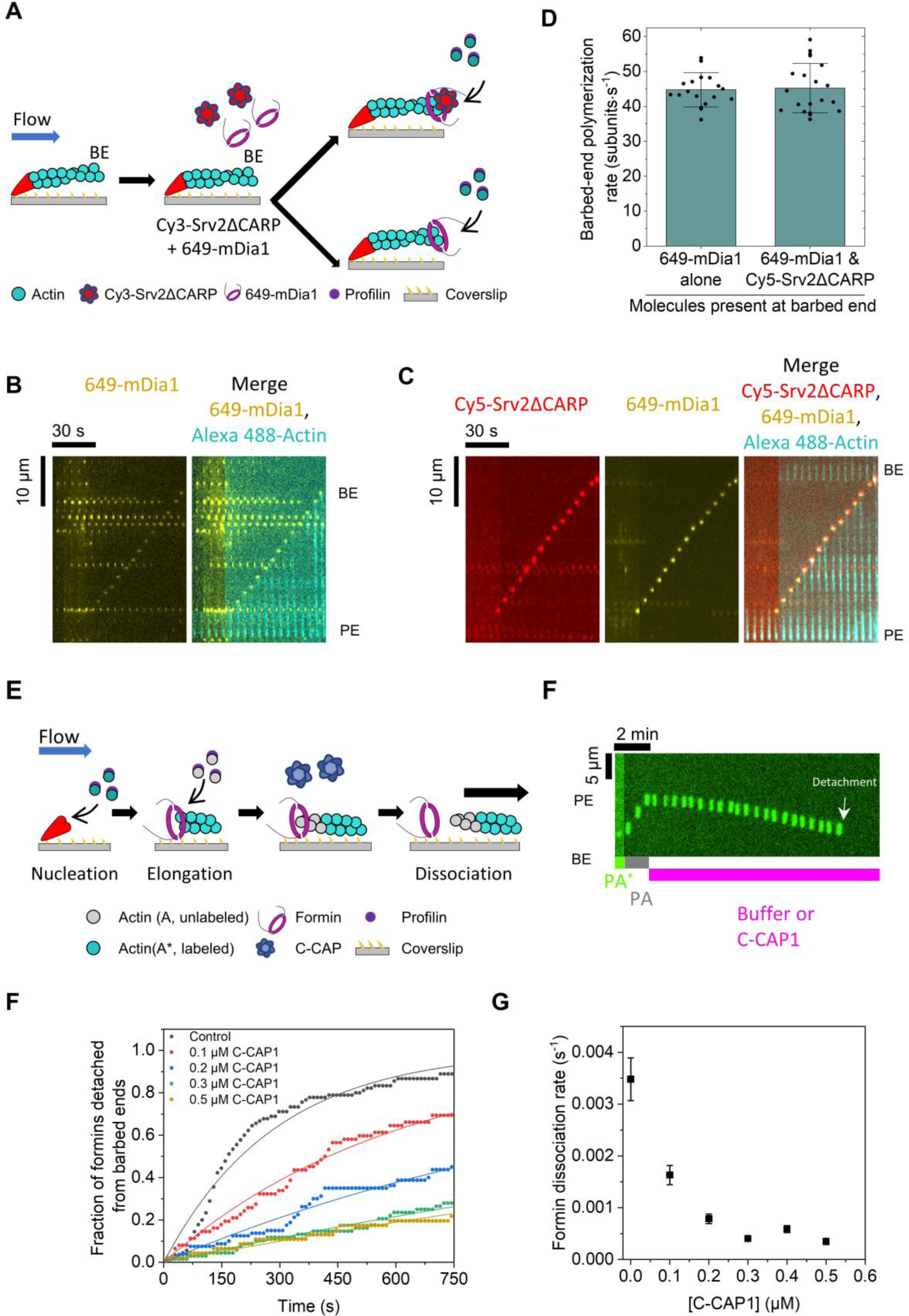
Srv2/CAP stabilizes barbed-end binding of formin. **(A)** Schematic representation of the experimental strategy. Actin filaments with free barbed ends were assembled similarly as in Fig. 1A and exposed to either 649-mDia1 alone or together with Cy3-Srv2ΔCARP, and then exposed to 1 µM Alexa-488 G-actin and 4 µM profilin. Red colored Srv2ΔCARP molecules represent labeled protein. **(B)** Representative kymograph of an Alexa-488-labeled actin filament (cyan) bound to 649-mDia1 (yellow) at the barbed end. **(C)** Representative kymograph of an Alexa-488-labeled actin filament (cyan) bound to 649-mDia1 (yellow) and Cy3-Srv2ΔCARP (red) at the barbed end **(D)** Rates (± sd) of barbed-end elongation of 649-mDia-bound or 649-mDia- and Cy3-Srv2ΔCARP-bound barbed ends elongating from 1 µM Alexa-488 G-actin and 4 µM profilin. Number of filaments analyzed for each condition: 20. **(E)** Schematic representation of the experimental strategy. Actin filaments were nucleated from coverslip-anchored formins by introducing a flow containing 1 µM G-actin (15% Alexa-488 labeled) and 0.5 µM profilin. The filaments were then allowed to elongate in presence of 1 µM unlabeled G-actin and 4 µM profilin to ensure insertional elongation between fluorescent fragment and surface-anchored formins. These filaments were then exposed to a flow containing C-CAP in absence of profilin-actin. The survival fraction of filaments attached to formins was monitored as a function of time. Blue colored C-CAP molecules represent unlabeled protein. **(F)** Representative kymograph of a formin-anchored filament elongating from unlabeled monomers (see Fig. 3D) and then being exposed to buffer. Time point of filament detachment from formin is indicated. **(G)** Fraction of filaments attached to formin as a function of time in presence of varying concentrations of C-CAP. Experimental data (symbols) are fitted to a single-exponential function (lines) to determine formin dissociation rate. Number of filaments analyzed for each condition (from low to high C-CAP concentration): 90, 62, 80, 68, 46) **(H)** Formin dissociation rate as a function of C-CAP concentration, determined from data shown in (G).

We then asked if Srv2/CAP altered barbed-end processivity of formin. Since processivity is influenced by actin labelling fraction, elongation rate and presence of profilin [42], we employed an alternative strategy in (absence of profilin and G-actin) which we and others have previously used to study processivity of formin (Fig 3E) [38, 40, 42]. Actin filaments were nucleated by exposing coverslip-anchored formins to a solution containing fluorescent actin monomers and profilin (Fig. 3E). To ensure that insertional polymerization was occurring at anchored formins, a solution containing profilin and unlabeled actin monomers was introduced. As a result, fluorescent segments appeared to move along the flow as filaments elongated (Fig, 3E,F). These actin filaments were then exposed to a solution containing a range of C-CAP concentrations (in absence of profilin-actin). The time-dependent disappearance of filaments from the field of view (due to their dissociation from surface-anchored formins) was analyzed to determine formin’s dissociation rate from barbed ends. We found that formins remained bound to filaments for longer durations in the presence of C-CAP (Fig. 3F). Formin’s dissociation rate decreased with increasing C-CAP concentration. At saturating C-CAP concentration, we observed a 10-fold increase in barbed end processivity of formin (Fig. 3G). Consistent with single-molecule analysis, C-CAP did not alter the depolymerization rate of formin-bound barbed ends. Together, our data suggests that while C-CAP stabilizes formin’s association at barbed ends, it does not alter formin’s effects on the polymerization/depolymerization rate.

### Srv2/CAP colocalizes with CP at barbed ends and accelerates uncapping

We then sought to investigate if CAP might also alter CP’s barbed-end binding. A SNAP-tagged construct of mouse capping protein (SNAP-CP) was expressed and labeled with benzylguanine functionalized green-excitable (549-CP) fluorescent dye. As seen previously, SNAP-tagging and labelling did not alter CP’s interactions with the barbed-end [40, 41]. Alexa-488 labeled actin filaments with free barbed ends were transiently exposed to 549-CP in presence of profilin and G-actin (Fig. 4A). Appearance of a 549-CP signal at barbed ends coincided with filaments switching from elongating to paused state. Upon subsequent exposure to a solution containing Cy5-Srv2ΔCARP and profilin-actin, we observed that Srv2ΔCARP colocalized with 549-CP at barbed ends (Fig. 4B). During co-localization, filaments remained paused; no change in filament length was observed. We then asked if Srv2/CAP might influence barbed-end residence of CP? We employed a strategy similar to one used for investigating formins’ processivity. Preformed fluorescently-labeled actin filaments were captured by coverslip-anchored CP in the mf-TIRF chamber (Fig. 4C). Time-dependent disappearance of filaments from the field of view was recorded and used to determine the uncapping rate. In presence of C-CAP, the uncapping rate increased linearly with C-CAP concentration (Fig D-G). Notably, although we only observed a maximum of 4-fold enhancement in CP’s dissociation rate from the barbed end, given the linear nature of concentration-dependence we expect further enhancement at higher C-CAP concentrations. Our results add CAP to the list of proteins with uncapping abilities namely formin [38], CARMIL [43], twinfilin [44], cofilin [35], VopF [45].

**Fig. 4:**
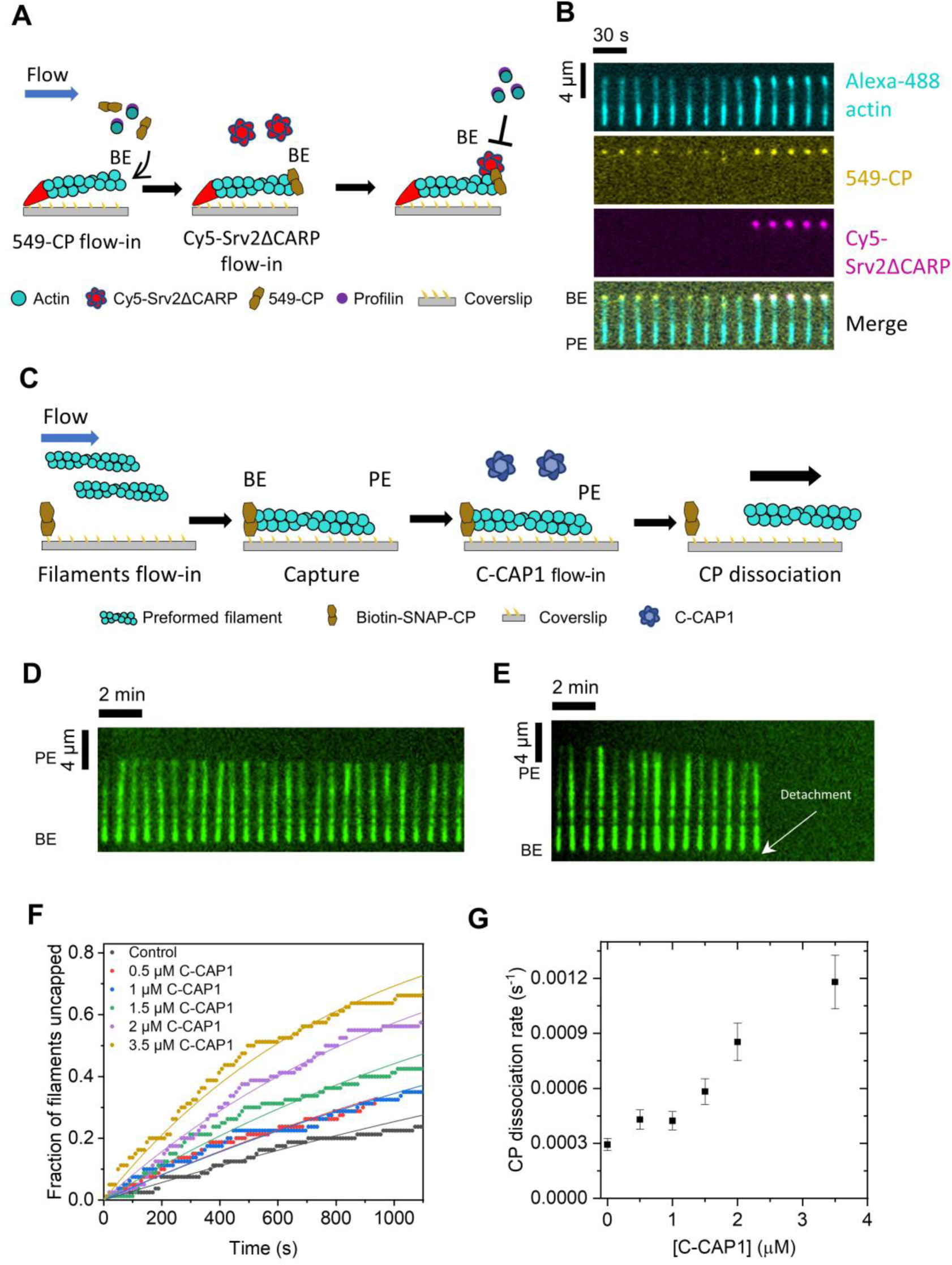
Srv2/CAP co-localizes with CP at barbed ends and promotes uncapping. **(A)** Schematic representation of the experimental strategy. Actin filaments with free barbed ends were assembled similarly as in Fig. 1A and were transiently exposed to 10 nM 549-CP in presence of 1 µM Alexa-488 G-actin and 4 µM profilin. The filaments were then exposed to 100 nM Cy5-Srv2ΔCARP. Red colored Srv2ΔCARP molecules in the schematic represent labeled protein. **(B)** Representative kymograph of an Alexa-488-labeled actin filament with 549-CP and Cy5-Srv2ΔCARP simultaneously bound to the barbed end. **(C)** Schematic representation of the experimental strategy. Pre-formed Alexa-488 labelled ADP-actin filaments were captured by coverslip-anchored biotinylated SNAP-CP anchored on the glass coverslip. Varying concentrations of C-CAP were then introduced into the chamber, and filament detachment was monitored. BE, barbed end; PE, pointed end. Blue colored C-CAP molecules represent unlabeled protein. **(D)** Representative kymograph of a CP-anchored filament in buffer **(E)** Same as (D) but with 3 µM C-CAP **(F)** Fraction of filaments attached to CP as a function of time in presence of varying concentrations of C-CAP. Experimental data (symbols) are fitted to a single-exponential function (lines) to determine CP dissociation rate. Number of filaments analyzed for each condition: 80. **(G)** CP dissociation rate as a function of C-CAP concentration, determined from data shown in (F).

### Role of WH2 domains in barbed end effects of CAP

WH2 domains have primarily been considered as actin monomer-binding motifs which promote nucleation [22, 46]. WH2-containing nucleators include SPIRE [47], Cordon Bleu [48], Leiomodin [49], VopF/VopL [50, 51], and Sca2 [52, 53]. WH2 binding to barbed ends is considered less favorable due to the flatter F-actin conformation. How then might WH2 domain of CAP interact with the barbed end? There are two possible lines of reasoning. First, although the WH2-binding site is largely obscured on subunits in the filament, the binding site is fully exposed at the barbed end [46]. Second, a high-resolution barbed-end structure recently published in a landmark study by the Dominguez lab showed that the W loop and the C-terminus of terminal actin’s hydrophobic cleft undergoes a G-actin-like conformation at the barbed end [54]. These conformational changes likely play a role in barbed-end effects of WH2-containing proteins like CAP, N-WASP [55], WASP [56] and VopF [45]. To our knowledge, ours is the first report of WH2 domains facilitating processive barbed-end depolymerization.

In addition to binding free barbed and pointed ends, we find that CAP can also bind formin-bound and CP-bound barbed ends. Co-localization of CAP with either of these two proteins does not alter their respective individual effects on barbed-end elongation. Similar observations were previously made for budding yeast twinfilin i.e., presence of yeast Srv2 increased the processivity but not barbed-end depolymerization rate of twinfilin [36]. Interestingly, the increase in processivity of twinfilin was mediated by the N-terminal half of Srv2 and not the C-terminal half! Thus, CAP alters barbed-end residence time of elongators (formin), blockers (capping protein) and depolymerases (twinfilin). In future, it will be interesting to investigate if CAP might also co-localize with formin-CP-twinfilin complex and influence the dynamics of this multiprotein barbed-end complex [40].

We believe our results bring new insights into how multiple proteins might co-exist at the barbed end. CP binds the barbed end primarily via interactions of its β- and α-tentacle with hydrophobic cleft of the terminal and penultimate actin subunits [54, 57, 58]. WH2 domains also target the same hydrophobic cleft on actin [59]. Deletion of β-tentacle “only” weakens barbed-end affinity of CP by 300-fold in comparison to 5000-fold weakening in absence of α -tentacle [58]. We speculate that destabilization of CP by CAP is primarily caused by a competition between the β-tentacle and the WH2 domain as reported recently [59]. This is likely further aided by splaying of the barbed-end subunits due to CP binding [54]. We believe these structural observations open the possibility for a plausible mechanism for destabilization of CP by WH2 domains, and allowing for faster uncapping by CAP. Another WH2 domain protein VopF has also been previously reported to displace CP from barbed ends [45].

While our manuscript was under preparation, we became aware of another study with results that partially overlap with our findings [18]. Although effects of CAP on barbed end depolymerization are qualitatively consistent between the studies, ADP-actin depolymerization rates were 4-fold faster rate in our study (∼58 su/s vs 15 su/s). We believe this disparity originates from differences in the filament aging process. While we aged filaments for 15 minutes after polymerization to explicitly convert them to ADP-actin, filaments in the other study were immediately exposed to CAP following their polymerization to mimic *in vivo* conditions. As a result, the filaments likely contained a mix of ADP and ADP-P_i_ subunits. Further, Alimov et al also report that CARP domains are required for barbed-end depolymerization. They found that a modified C-CAP construct consisting of poly-proline regions, WH2 motif and a GST tag but not the CARP domain failed to promote depolymerization. In contrast, we found that Srv2ΔCARP which contained the WH2 domain but lacked the CARP domain depolymerized barbed ends as fast as C-Srv2 which contained both WH2 and CARP. Additionally, the authors found that C-CAP promotes dissociation of formin from barbed ends. In contrast, we observe that C-CAP increased barbed-end residence time of formin by 4-fold. We further note that co-localization with CAP did not change the elongation rate of formin in our experiments.

### Concluding remarks and cellular implications

Combining our observations with previously known activities of CAP, it depolymerizes barbed ends, accelerates cofilin-mediated pointed end depolymerization [6, 7], uncaps CP, stabilizes barbed-end binding of formin and twinfilin [36], collaborates with Abp1 to bundle filaments [12], and promotes nucleotide exchange [13–15]. In addition, a complex of lysine-acetylated actin and CAP regulates autoinhibition of formin INF2 [60]. In light of the wide diversity of effects CAP exerts on actin dynamics directly and via other proteins, it would not be an exaggeration to call CAP the “swiss army knife” of actin dynamics!

The phenomenon of actin “treadmilling” has formed the central bedrock of our understanding of actin dynamics for over three decades [1, 28]. “Treadmilling” entails actin filaments polymerizing at their barbed ends and depolymerizing from their pointed ends. Our discovery of barbed-end depolymerization by CAP together with previously reported barbed-end effects of twinfilin [32, 36] and pointed-end polymerization by VopF [61] further challenge the universality of actin “treadmilling”.

Although actin regulatory proteins have classically been studied one at a time, we now understand that in cells, they act simultaneously in multiprotein teams, often at the same site on a filament. Our results shed light on novel multicomponent activities of CAP at the barbed end i.e., promoting uncapping and stabilizing formin. It was previously shown that when present alone, CP promotes displacement of formin from the barbed end [38, 41]. However, addition of the uncapper twinfilin promoted formin-mediated actin assembly [40]. Since CAP increases twinfilin’s processivity [36] and uncaps CP[44], we speculate that in the complex intracellular milieu, CAP and twinfilin might together act as a highly potent uncapper and in turn promote formin-mediated assembly. Nevertheless, we are fully aware that future cellular studies are needed to test the physiological relevance of *in vitro* mechanisms and predictions reported here. We also believe that future Cryo-EM structural studies will be key to gaining deeper mechanistic insights on how CAP interacts with barbed ends, alone and together with formin and CP.

## Acknowledgements

We are grateful to Pekka Lappalainen for generously sharing CAP plasmids. We thank Shoichiro Ono, Ankita and Heidi Ulrichs for critical reading of the manuscript and for valuable feedback. We thank all Shekhar lab members, and especially Heidi Ulrichs, for help with protein purification. This work was supported by NIH NIGMS grant R35GM143050 to S.S. and startup funds from Emory University to S.S.

## Author contributions

EMT conducted experiments and analyzed data. EMT and SS prepared figures. SS designed experiments and supervised the project. SS wrote the manuscript and both authors contributed to the editing. SS acquired funding.

## Methods

### Purification and labeling of actin

Rabbit skeletal muscle actin was purified from acetone powder generated from frozen ground hind leg muscle tissue of young rabbits (PelFreez, USA). Lyophilized acetone powder stored at −80°C was mechanically sheared in a coffee grinder, resuspended in G-buffer (5 mM Tris-HCl pH 7.5, 0.5 mM Dithiothreitol (DTT), 0.2 mM ATP and 0.1 mM CaCl_2_), and cleared by centrifugation for 20 min at 50,000 × *g*. Supernatant was collected and further filtered with Whatman paper. Actin was then polymerized overnight at 4°C, slowly stirring, by the addition of 2 mM MgCl_2_ and 50 mM NaCl to the filtrate. The next morning, NaCl powder was added to a final concentration of 0.6 M and stirring was continued for another 30 min at 4°C. Then, F-actin was pelleted by centrifugation for 150 min at 280,000 × *g*, the pellet was solubilized by dounce homogenization and dialyzed against G-buffer for 48 h at 4°C. Monomeric actin was then precleared at 435,000 × *g* and loaded onto a Sephacryl S-200 16/60 gel-filtration column (Cytiva, USA) equilibrated in G-Buffer. Fractions containing actin were stored at 4°C.

To fluorescently label actin, G-actin was polymerized by dialyzing overnight against modified F-buffer (20 mM PIPES pH 6.9, 0.2 mM CaCl_2,_ 0.2 mM ATP, 100 mM KCl)[62]. F-actin was incubated for 2 h at room temperature with a 5-fold molar excess of Alexa-488 NHS ester dye (Thermo Fisher Scientific, USA). F-actin was then pelleted by centrifugation at 450,000 × *g* for 40 min at room temperature, and the pellet was resuspended in G-buffer, homogenized with a dounce and incubated on ice for 2 h to depolymerize the filaments. The monomeric actin was then re-polymerized on ice for 1 h by addition of 100 mM KCl and 1 mM MgCl_2_. F-actin was once again pelleted by centrifugation for 40 min at 450,000 × *g* at 4°C. The pellet was homogenized with a dounce and dialyzed overnight at 4°C against 1 L of G-buffer. The solution was precleared by centrifugation at 450,000 × *g* for 40 min at 4°C. The supernatant was collected, and the concentration and labeling efficiency of actin was determined.

### Purification of profilin

Human profilin-1 was expressed in *E. coli* strain BL21 (pRare) to log phase in LB broth at 37°C and induced with 1 mM IPTG for 3 h at 37°C. Cells were then harvested by centrifugation at 15,000 × *g* at 4°C and stored at -80°C. For purification, pellets were thawed and resuspended in 30 mL lysis buffer (50 mM Tris-HCl pH 8, 1 mM DTT, 1 mM PMSF protease inhibitors (0.5 μM each of pepstatin A, antipain, leupeptin, aprotinin, and chymostatin)) was added, and the solution was sonicated on ice by a tip sonicator. The lysate was centrifuged for 45 min at 120,000 × *g* at 4°C. The supernatant was then passed over 20 ml of poly-L-proline conjugated beads in a disposable column (Bio-Rad, USA). The beads were first washed at room temperature in wash buffer (10 mM Tris pH 8, 150 mM NaCl, 1 mM EDTA and 1 mM DTT) and then washed again with 2 column volumes of 10 mM Tris pH 8, 150 mM NaCl, 1 mM EDTA, 1 mM DTT and 3 M urea. Protein was then eluted with 5 column volumes of 10 mM Tris pH 8, 150 mM NaCl, 1 mM EDTA, 1 mM DTT and 8 M urea. Pooled and concentrated fractions were then dialyzed in 4 L of 2 mM Tris pH 8, 0.2 mM EGTA, 1 mM DTT, and 0.01% NaN_3_ (dialysis buffer) for 4 h at 4°C. The dialysis buffer was replaced with fresh 4 L buffer and the dialysis was continued overnight at 4°C. The protein was centrifuged for 45 min at 450,000 × *g* at 4°C, concentrated, aliquoted, flash frozen in liquid N_2_ and stored at -80°C.

### Purification of Cofilin-1

Human Cofilin-1 was expressed in *E.coli* BL21 DE3 cells. Cells were grown in Terrific Broth to log phase at 37⁰C, and then expression was induced overnight at 18⁰C by addition of 1 mM IPTG. Cells were collected by centrifugation and pellets were stored at -80⁰C. Frozen pellets were thawed and resuspended in lysis buffer (20 mM Tris pH 8.0, 50 mM NaCl, 1 mM DTT, and protease inhibitors (0.5 µM each of pepstatin A, antipain, leupeptin, aprotinin, and chymostatin). Cells were lysed with a tip sonicator while being kept on ice. The cell lysate was centrifuged at 150,000 × *g* for 30 min at 4⁰C. The supernatant was loaded on a 1 ml HisTrap HP Q column (GE Healthcare, Pittsburgh, PA), and the flow-through was collected and dialyzed against 20 mM HEPES pH 6.8, 25 mM NaCl, and 1 mM DTT. The dialyzed solution was then loaded on a 1 ml HisTrap SP FF column (GE Healthcare, Pittsburgh, PA) and eluted using a linear gradient of NaCl (20-500 mM). Fractions containing protein were concentrated, dialyzed against 20 mM Tris pH 8.0, 50 mM KCl, and 1 mM DTT, flash frozen in liquid N_2_ and stored at -80⁰C.

### Purification and biotinylation of SNAP-CP

SNAP-CP [41] was expressed in E. coli BL21 DE3 by growing cells to log phase at 37°C in TB medium, then inducing expression using 1 mM IPTG at 18°C overnight. Cells were harvested by centrifugation and pellets were stored at −80°C. Frozen pellets were resuspended in lysis buffer (20 mM NaPO_4_ pH 7.8, 300 mM NaCl, 1 mM DTT, 15 mM imidazole, 1 mM PMSF) supplemented with a protease inhibitor cocktail (0.5 µM each of pepstatin A, antipain, leupeptin, aprotinin, and chymostatin). Cells were lysed by sonication with a tip sonicator while keeping the tubes on ice. The lysate was cleared by centrifugation at 150,000 x g for 30 min at 4°C. The supernatant was then flowed through a HisTrap column connected to a Fast Protein Liquid Chromatography (FPLC) system. The column with the bound protein was first extensively washed with the washing buffer (20 mM NaPO_4_ pH 7.8, 300 mM NaCl, 1 mM DTT and 15 mM imidazole) to remove non-specifically bound proteins. SNAP-CP was then eluted with 250 mM imidazole in 20 mM NaPO_4_ pH7.8, 300 mM NaCl, and 1 mM DTT. The eluted protein was concentrated and labelled either with Benzylguanine Biotin or SNAP-surface-549 (New England Labs) according to the manufacturer’s instructions. Free biotin (or dye) was removed using size-exclusion chromatography by loading the labelled protein on a Superose 6 gel filtration column (GE Healthcare, Pittsburgh, PA) eluted with 20 mM HEPES pH 7.5, 150 mM KCl, 0.5 mM DTT. Fractions containing the protein were pooled, aliquoted, snap frozen in liquid N_2_ and stored at - 80°C.

### Purification, labeling and biotinylation of formin mDia1

Mouse his-tagged SNAP-mDia1 (FH1-FH2-C) formin was expressed in *E. coli*; BL21(DE3) pLysS cells. Cells were grown in Terrific Broth to log phase at 37°C. Expression was induced overnight at 18°C by addition of 1 mM IPTG. Cells were harvested by centrifugation at 11,200 × *g* for 15 min and the cell pellets were stored at -80°C. For purification, frozen pellets were thawed and resuspended in 35 mL lysis buffer (50 mM sodium phosphate buffer pH 8, 20 mM imidazole, 300 mM NaCl, 1 mM DTT, 1 mM PMSF and protease inhibitors (0.5 μM each of pepstatin A, antipain, leupeptin, aprotinin, and chymostatin)). Cells were lysed using a tip sonicator while being kept on ice. The cell lysate was then centrifuged at 120,000 × *g* for 45 min at 4°C. The supernatant was then incubated with 1 mL of Ni-NTA beads (Qiagen, USA) while rotating for 2 h at 4°C. The beads were then washed three times with the wash buffer (50 mM sodium phosphate buffer pH 8, 300 mM NaCl, 20 mM imidazole and 1 mM DTT) and were then transferred to a disposable column (Bio-Rad, USA). Protein was eluted using the elution buffer (50 mM phosphate buffer pH 8, 300 mM NaCl, 250 mM imidazole and 1 mM DTT). Fractions containing the protein were concentrated and loaded onto a size exclusion Superdex 200 increase 10/300 GL column (Cytiva, USA) pre-equilibrated with 20 mM HEPES pH 7.5, 150 mM KCl, 10% glycerol and 0.5 mM DTT. The eluted protein was concentrated and labelled either with Benzylguanine Biotin or SNAP-surface-649 (New England Labs) according to the manufacturer’s instructions. Free biotin (or dye) was removed using a Superdex 200 increase 10/300 GL column (Cytiva, USA). Fractions containing the protein were pooled, aliquoted, snap frozen in liquid N_2_ and stored at -80°C.

### Purification of CAP1 and Srv2 peptides

Mouse CAP1 and *S. cerevisiae* Srv2 peptides were expressed as His-tagged constructs in E. coli BL21 pRARE or pLysS by growing cells to log phase at 37°C in TB medium. Full-length CAP1 and C-CAP contained additional SUMO and 3C cleavage sites respectively [7]. Cells were induced with 1 mM IPTG at 18°C overnight. Cells were harvested by centrifugation and pellets were stored at −80°C. Frozen pellets were resuspended in 50 mM NaPO_4_ pH 8.0, 1 mM PMSF, 1 mM DTT, 20 mM imidazole, 300 mM NaCl and protease inhibitors as described above. Cells were lysed by sonication with a tip sonicator while keeping the tubes on ice. The lysate was cleared by centrifugation at 150,000 x g for 30 min at 4°C. The lysate was incubated for two hours with Ni-NTA beads (Qiagen). Non-specifically bound proteins were removed by washing the beads with 20 mM NaPO_4_ pH 8.0, 50 mM imidazole, 300 mM NaCl and 1 mM DTT. The bound protein was then eluted using 250 mM imidazole in the same buffer. Fractions containing the protein were concentrated and then further purified by size-exclusion chromatography using Superdex 75 Increase or Superdex 200 Increase gel-filtration columns (Cytiva, USA) equilibrated in 5 mM HEPES pH7.4, 100 mM NaCl and 1 mM DTT. Peak fractions were pooled, the protein was aliquoted, snap frozen in liquid N_2_ and stored at -80 °C.

For FL-CAP1 and C-CAP1, the His-tag was cleaved prior to gel-filtration. The eluted protein from Ni-NTA beads was first dialyzed against 20 mM HEPES pH 7.4, 300 mM NaCl and 2 mM DTT, and then incubated overnight either with SUMO protease (Sigma Aldrich) or PreScission Protease. The cleaved tags were removed by incubation with Ni-NTA and/or GST beads.

### Labeling of Srv2ΔCARP

For fluorescent labeling of Srv2ΔCARP, the same procedure as above was followed for unlabeled Srv2ΔCARP with the exception that 1 mM DTT in the elution buffer was replaced with 0.2 mM Tris(2-carboxyethyl)phosphine (TCEP) [63]. The eluted fractions were concentrated and incubated with at least fivefold molar excess of Cy3- or Cy5-maleimide dye (GE Healthcare, Pittsburgh, PA) for 30 min at 25 °C and additionally for 14 h at 4 °C. The excess dye was then quenched by addition of 5 mM DTT. Free dye was then separated from labeled protein using a PD-10 column with 10 mM imidazole pH 8, 50 mM KCl, 1 mM DTT and 5% glycerol. Labeled protein was concentrated. The protein was then aliquoted, snap frozen in liquid N_2_ and stored at −80 °C.

### Microfluidics-assisted TIRF (mf-TIRF) microscopy

Actin filaments were assembled in microfluidics-assisted TIRF (mf-TIRF) flow cells [62]. For all experiments, coverslips were first cleaned by sonication in Micro90 detergent for 20 min, followed by successive 20 min sonications in 1 M KOH, 1 M HCl and 200 proof ethanol for 20 min each. Washed coverslips were then stored in fresh 200 proof ethanol. Coverslips were then washed extensively with H_2_O and dried in an N_2_ stream. These dried coverslips were coated with 2 mg/mL methoxy-poly (ethylene glycol) (mPEG)-silane MW 2,000 and 2 µg/mL biotin-PEG-silane MW 3,400 (Laysan Bio, USA) in 80% ethanol (pH 2.0) and incubated overnight at 70°C. A 40 µm high PDMS mold with 3 of 4 inlets and 1 outlet was mechanically clamped onto a PEG-Silane coated coverslip. The chamber was then connected to a Maesflo microfluidic flow-control system (Fluigent, France), rinsed with TIRF buffer (10 mM imidazole pH 7.4, 50 mM KCl, 1 mM MgCl_2_, 1 mM EGTA, 0.2 mM ATP, 10 mM DTT, 1 mM DABCO) and incubated with 1% BSA and 10 µg/mL streptavidin in 20 mM HEPES pH 7.5, and 50 mM KCl for 5 min. Depending upon the needs of specific experiments, spectrin-seeds, biotinynlated-capping protein or biotinylated-formin was then anchored on the surface by flowing them in TIRF buffer for 5 min.

For ADP-P_i_ experiments (Fig. 1G), the filaments were maintained in modified TIRF buffer (regular TIRF buffer supplemented with inorganic phosphate : 10 mM imidazole pH 7.4, 34.8 mM K_2_HPO_4_ and 15.2 mM KH_2_PO_4_, 1 mM MgCl_2_, 1 mM EGTA, 0.2 mM ATP, 10 mM DTT, 1 mM DABCO) throughout the experiment.

### Image acquisition and analysis

Multi-wavelength time-lapse TIRF imaging was performed on a Nikon-Ti2000 inverted microscope equipped with a 40 mW 488 nm, 561 nm and 640 nm Argon lasers, a 60X TIRF-objective with a numerical aperture of 1.49 (Nikon Instruments Inc., USA) and an IXON LIFE 888 EMCCD camera (Andor Ixon, UK). One pixel was equivalent to 144 × 144 nm. Focus was maintained by the Perfect Focus system (Nikon Instruments Inc., Japan). Time-lapsed images were acquired using Nikon Elements imaging software (Nikon Instruments Inc., Japan).

Images were analyzed in Fiji [64]. Background subtraction was conducted using the rolling ball background subtraction algorithm (ball radius 5 pixels). Time-lased images were corrected for drift using Fiji Image Stabilizer plugin. For each condition, filaments were acquired across multiple fields of view. To determine the rate of depolymerization, the in-built kymograph plugin was used to draw kymographs of individual filaments. The kymograph slope was used to calculate barbed- or pointed-end depolymerization rate of each individual filament (assuming one actin subunit contributes 2.7 nm to filament length). Data analysis and curve fitting were carried out in Microcal Origin. All experiments were repeated at least three times and yielded similar results. Unless otherwise mentioned, the data shown are from one trial.

### Data availability

Data supporting the findings of this manuscript are available from the corresponding author upon reasonable request.

